# Identification and Validation of Mitophagy-Related Genes in Diabetic Retinopathy

**DOI:** 10.1101/2024.09.10.612286

**Authors:** Wenxuan Peng, Yulin Zou

## Abstract

**Background:** Diabetic retinopathy is one of the common chronic complications of diabetes, characterized by retinal microvascular and neurodegenerative impairment, and it is the primary cause of vision impairment and blindness in adults. Many studies have demonstrated that mitophagy plays a significant role in the pathological mechanism of DR. however, its mechanism is not yet fully clear and requires further research.

**Methods:** We obtained relevant datasets of diabetic retinopathy from the GEO database and used R language to screen for differentially expressed genes. We intersected these genes with mitophagy-related genes and identified differentially expressed mitophagy-related genes. We performed GO and KEGG analysis on the differentially expressed mitophagy-related genes, followed by PPI network analysis. Using Cytoscape software, we selected mitophagy hub genes. Finally, we further validated the expression of the mitophagy hub genes in an in vitro cell culture high-glucose model using quantitative real-time polymerase chain reaction (qRT-PCR).

**Results:** We identified 27 differentially expressed genes related to mitophagy by using R language, with 10 genes upregulated and 17 genes down regulated. We performed GO and KEGG enrichment analysis using R software to further study the potential biological functions of differentially expressed genes. Through PPI network analysis and Cytoscape software, we selected 10 hub genes associated with mitophagy. Finally, through qRT-PCR validation of these 10 hub genes, we found that the mRNA expression differences of MFN1, BNIP3L, GABARAPL1, and PINK1 genes were consistent with our bioinformatics analysis results.

**Conclusion:** We consider that MFN1, BNIP3L, GABARAPL1, and PINK1 may serve as potential biomarkers for diabetic retinopathy. The upregulation and downregulation of these genes provide new insights for further exploration of the role of mitophagy in the pathological mechanism of diabetic retinopathy. These genes can serve as new potential therapeutic targets for the treatment of diabetic retinopathy.

## 1. Introduction

Diabetic retinopathy is a series of diseases caused by the damage of retinal microvasculature due to diabetes.According to the World Health Organization (WHO), diabetic retinopathy accounts for 4.8% of the global cases of blindness (37 million)^[1]^.Diabetic retinopathy can be classified into different stages based on the degree and extent of the lesions, including mild non-proliferative diabetic retinopathy, moderate to severe non-proliferative diabetic retinopathy, and proliferative diabetic retinopathy.The development of DR is a complex process, and the known pathological mechanisms of DR are mainly due to oxidative stress induced by sustained high blood sugar, abnormal vascular permeability, microaneurysms, vascular occlusion, and dysfunction of endothelial cells. This leads to increased retinal vascular permeability, neurodegenerative changes, neovascularization, ultimately resulting in the breakdown of the blood-retinal barrier^[2-5]^.However, the pathology and mechanisms of diabetic retinopathy are still not fully understood.

Mitophagy, also known as mitochondrial autophagy or mitochondrial clearance, is a cellular self-regulation process that maintains the health and functionality of cells by degrading and eliminating damaged or aging mitochondria^[6]^. In mitophagy, cells engulf mitochondria by forming double-membrane structures called autophagosomes and transport them to lysosomes for degradation. This process helps to remove dysfunctional mitochondria, reduce oxidative stress, maintain energy supply balance, and promote cellular metabolic adaptation. Mitophagy plays an important role in maintaining cellular homeostasis, coping with environmental stress, and promoting healthy aging^[7]^.Many studies believed that mitophagy can prevent cells from apoptosis and alleviate toxicity in stressful environments^[8]^.Studies have shown that dysfunction of mitophagy in Müller cells under high glucose conditions plays a crucial role in the pathogenesis of diabetic retinopathy (DR).In high-glucose environment, Müller cells exhibit impaired mitophagy, leading to the accumulation of autophagosomes in the cytoplasm and resulting in the release of large amounts of VEGF^[9]^.Some studies suggest that in high-glucose environment, PINK1/PARKIN-mediated mitophagy can promote the degradation of damaged mitochondria, thereby promoting cell survival.^[10]^.

Therefore, this study aimed to identify differentially expressed genes (DEGs) in the retina of patients with diabetic retinopathy (DR) and normal retinas. These DEGs were then intersected with the mitophagy dataset to first obtain differentially expressed genes related to mitophagy (DE-Mitophagy),we further performed functional enrichment analyses and Protein-protein interaction(PPI).We identified the top 10 genes in the PPI interaction network and designated them as hub genes.Finally, through qRT-PCR validation of these 10 hub genes in vitro DR model.Our findings revealed the role and mechanism of mitophagy in the progression of DR, new biomarkers.Targeted regulation of mitophagy can prevent the progression of DR (diabetic retinopathy), making it a potential therapeutic target for DR and providing more directions for its prevention and treatment.

## 2. Materials and methods

### 2.1. Data collection and preprocessing of mitophagy-related gene

We obtained the DR datasets used for this study from the GEO database.In this study, the GSE60436 mRNA Expression profiling by array dataset was downloaded from GEO (https://www.ncbi.nlm.nih.gov/geo/). GSE60436 database includes 6 samples from patients with diabetic retinopathy and 3 control samples from normal retinas and is based on GPL6884 (Illumina HumanWG-6 v3.0 expression beadchip Platform).

We used R version 4.3.1 to process the data and conducted gene ID conversion on the GSE60436 dataset and converted the Ensembl IDs to gene symbols and executed ID conversion in the GSE60436 dataset. The average expression value was deemed as the gene expression value when multiple Ensembl IDs/probes corresponded to the same gene symbol.

Genes related to mitophagy, called mitophagy-related genes (MRGs), were obtained from a public database(https://www.kegg.jp/entry/pathway+hsa04137).103 MRGs were identified and used for further analysis.The workflow of this study is shown in Figure 1.

**FIGURE 1.**
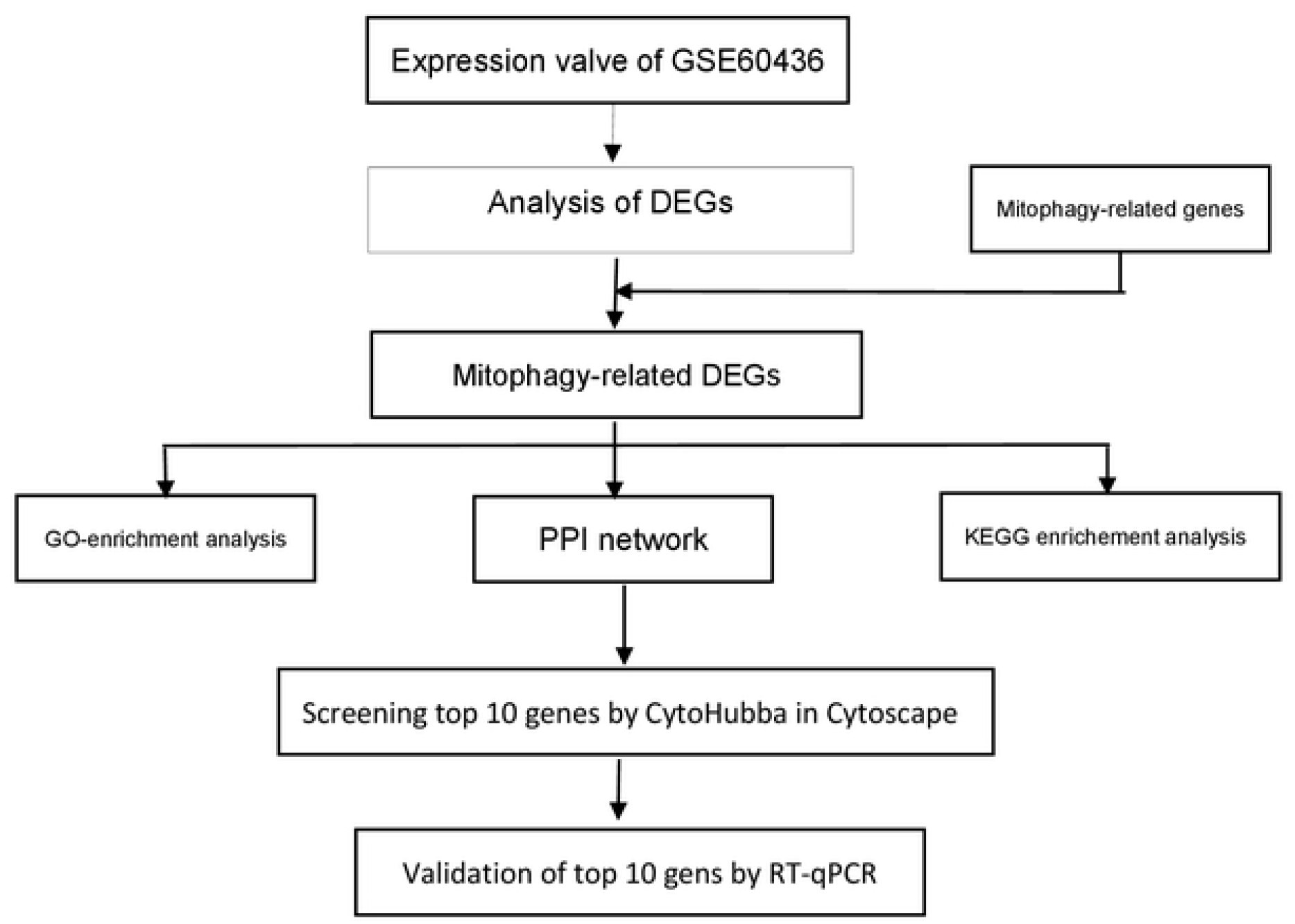
Experimental procedure diagram. The gene expression profiles of fibrovascular membranes extracted from the retinal of 3 non-diabetic individuals and 6 patients with DR in the GSE60436 dataset.103 mitophagy-related genes were collected from Kegg (https://www.kegg.jp/entry/pathway+hsaO4137). 10 genes up-regulated and 17 genes down-regulated in expression were screened by differential analysis. After enrichment analysis and PPI network analysis. 10 hub genes were determined. We further validated the in vitro diabetic retinopathy model through qRT-PCR. DEGs, differentially expressed genes; PPI, protein-protein interaction: qRT-PCR. quantitative real-time polymerase chain reaction.

### 2.2 Recognition of Mitophagy-related genes

We used R programming language.(version4.3.1;https://www.r-project.org/) and Bioconductor sofware package(http://www.bioconductor.org) to process and correct the raw data. GSE60436 mRNA Expression profile of normalization and quality control were performed by DESeq2 packages.Using the GEOquery and limma software packages to analyze the GSE60436 dataset and identify differentially expressed genes.We used Independent Principal Component Analysis (IPCA) to validate the repeatability of the GSE60436 database.The standard of statistical significance was |log2FC| > 0.5 and adjusted p < 0.05. MRGs were visualized in a volcano map based on the analysis.Finally, we created a volcano plot visualization using “umap” software package.In addition, we also constructed a Venn diagram and a heatmap for visual analysis of differentially expressed genes.

### 2.3 Functional enrichment analyses of Mitophagy-related genes DEGs

We utilized Gene Ontology (GO) and Kyoto Encyclopedia of Genes and Genomes (KEGG) enrichment analyses using the clusterProfiler for differentially expressed genes related to mitophagy, and visualize the results by generating a bar chart using the ggplot2 package for further analysis.

### 2.4. PPI construction network of DE-MRGs and hub gene identification

We utilized the STRING online database to construct the PPI interaction network and visualized it using Cytoscape software for further analysis.Based on the cytoHubba plugin in Cytoscape software, we identified the top 10 genes in the PPI interaction network and designated them as hub genes,which were computed base on the maximal clique centrality(MCC).

### 2.5. Cell Culture and Cell Grouping

We obtained human retinal capillary endothelial cells (HRCECs) from the Procell Life Science&Technology in China (CP-H130) and cultured in DMEM(Solarbio,CHINA) medium containing 10% fetal bovine serum (FBS) (Pricella, CHINA) and 1% Antibiotic-Antimycotic at 37°C with 5% carbon dioxide.

We implanted logarithmic growth phase cells with ideal development circumstances into six-well plates with 1.2×10^6^ cells per well.We divided the cells into two groups: the high glucose-treated group and the normal group.The high glucose-treated group was cultured in a medium containing 30mmol/L D-glucose, while the normal group was cultured in medium containing 5.5mmol/L D-glucose. Each well was supplemented with 2.5mL of the respective medium and incubated at 37°C for 48 hours.

### 2.6. RNA Extraction and qRT-PCR

We extracted RNA from human retinal capillary endothelial cells using TRIZOL(TransZol,CHINA),RNA reverse transcription was performed using the EasyScript®RT/RI Enzyme Mix(TransZol,CHINA) to synthesize cDNA,Finally, we performed qRT-PCR using the 2×Green qPCR SuperMix(TransZol,CHINA).We used primers synthesized at Shanghai Sangon Biotech Co., Ltd.We computed the mRNA expression quantification using the 2^-Δ Δ Ct method.We used β-actin as the reference gene mRNA.

### 2.7. Western Bolot

HUVECs were lysed with RIPA buffer and protein concentration was determined using the BCA protein assay. Each sample was normalized with loading buffer.

Subsequently, equal amounts of protein lysates were separated by 12% SDS-PAGE and transferred onto PVDF membranes at a constant current of 300 mA. PVDF membranes were blocked with 5% non-fat milk dissolved in TBST solution at room temperature for 2 hours. The membranes were then incubated overnight at 4°C with primary antibodies, followed by a 2-hour incubation at room temperature with secondary antibodies. After washing with TBST, the bands were visualized using an ECL detection system.

### 2.8. Statistical Analysis

We used GraphpadPrism software (version9.0.0) software to perform statistical analysis on the experimental data, and 3 independent experiments were executed. The gene expression level of the sample was matched by unpaired Student’s t-test, and the difference was considered to be statistically significant when P < 0.05.

## 3. RESULTS

### 3.1. Identification of DEGs and mitophagy-related DEGs

We obtained the Expression profiling by array dataset GSE60436 from the GEO database, and selected 3 non-diabetic individuals (Normal group) and 3 patients with FVMs(Fibrovascular membranes)with active NV proliferative diabetic retinopathy(PDR) and 3 patients FVMs with no active NV (PDR) (DR group)(Figures 2A).Next, we taken the intersection of the 6412 differentially expressed genes(logFC>0.5 and<-0.5 and adj.P.Val<0.05)and 103 mitophagy genes by R software.We obtained 27 differentially expressed genes, with 10 genes up-regulated and 17 genes down-regulated in expression. VennDiagram shows the intersection results(Figures 2B).and then graphed 27 differentially expressed mitophagy-related genes between the DR group and the normal group. A heatmap was used to display the gene expression levels of these differentially expressed genes related to mitophagy in the DR and normal samples.(Figures 3A)

**FIGURE 2.**
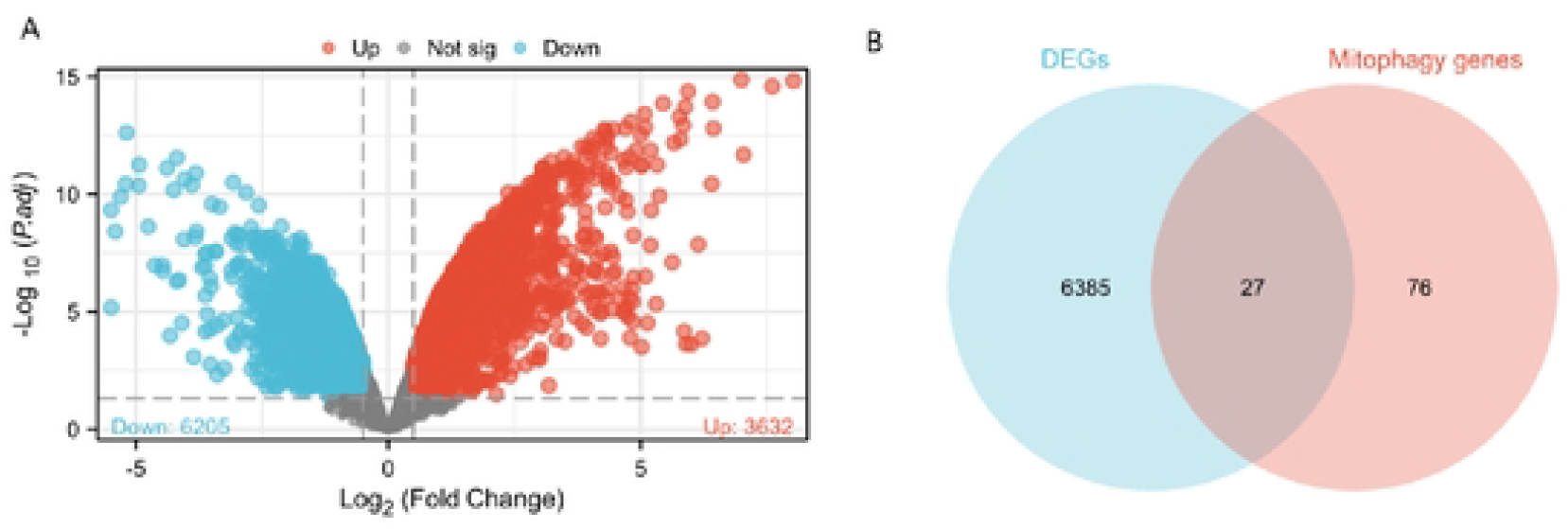
Differentially expressed mitophagy-related genes in DR patients (DR group) and non-diabetic individuals (Normal group). (A) Volcano plot of differentially expressed genes. The red dots in the picture represent significantly up-regulated genes, blue dots represent significantly down-regulated genes, grey dots represent genes that are not differentially expressed (B)Venn Diagram showed 6412 DEGs(Log2>0.5,P.adj<0.05) were intersected with the mitophagy dataset to obtain 27MEGs.

**FIGURE 3.**
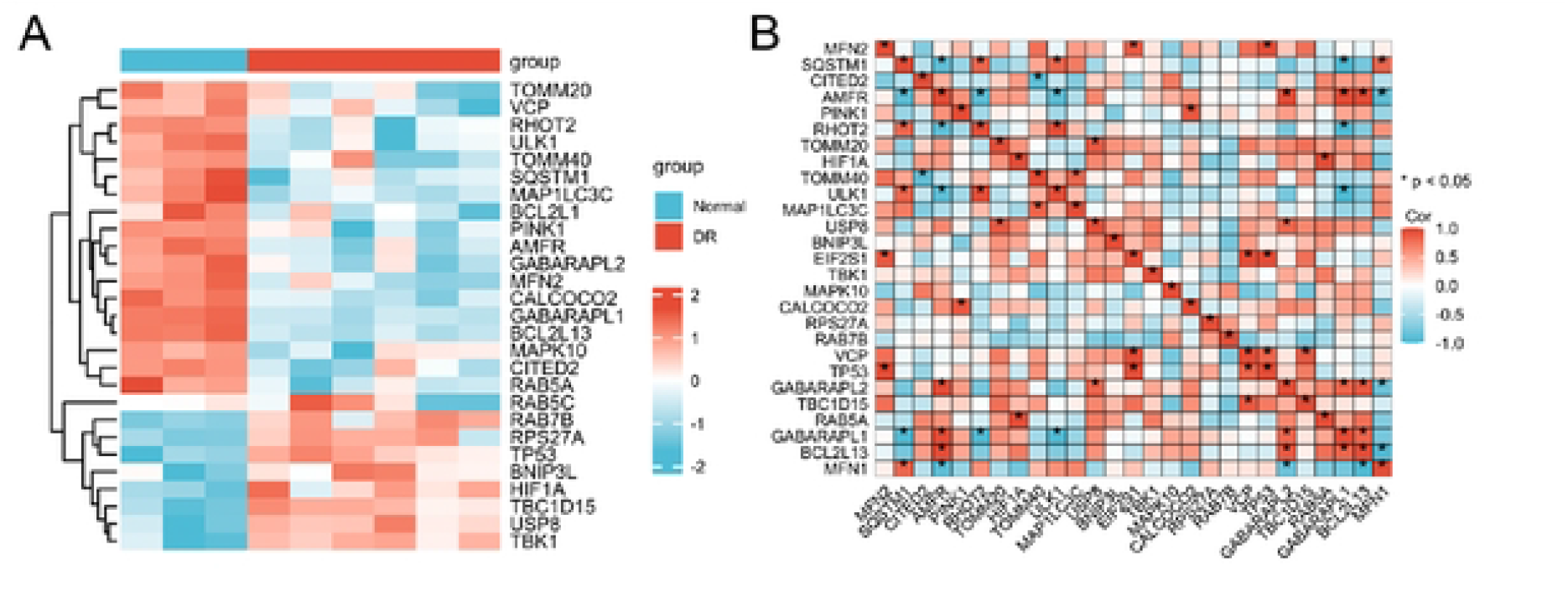
Correlation analysis of 27 differentially expressed mitophagy-related genes. (A), Correlation heatmap.

### 3.2. Correlation Expression of Differentially Expressed Mitophagy-Related Genes

We analyzed the expression correlation of 27 mitophagy-related genes using ggplot2 package to perform pairwise correlation analysis of genes and found a high degree of correlation among the genes.(Figures 3B)

### 3.3. Functional and Pathway Enrichment analyses of The Differentially Expressed Mitophagy-Related Genes

We performed GO and KEGG enrichment analysis using R software to further study the potential biological functions of differentially expressed genes. GO enrichment analysis displayed the most prominent projects involved regulation of mitophagy, positive regulation of catabolic process,macroautophagy, positive regulation of autophagy (BPs);outer membrane, mitochondrial outer membrane, organelle outer membrane (CCs); ubiquitin-like protein ligase binding ubiquitin protein ligase binding, GTP binding, GTPase activity (MFs)(Figures 4A–D).The bean plot shows expresion differences of 10 hub genes in dr and control groups. ROC curve of 10 hub genes showed that the diagnostic accuracies of MFN2,SQSTM1,PINK1,ULK1,BNIP3L,CALCOCO2,GABARAPL2,GABARAPL1, BCL2L13,MFN1 for DR were 100%,100%,100%,100%,94.4%,100%,100%,100%,100%,100%.(Figures 5A-C)

**FIGURE 4.**
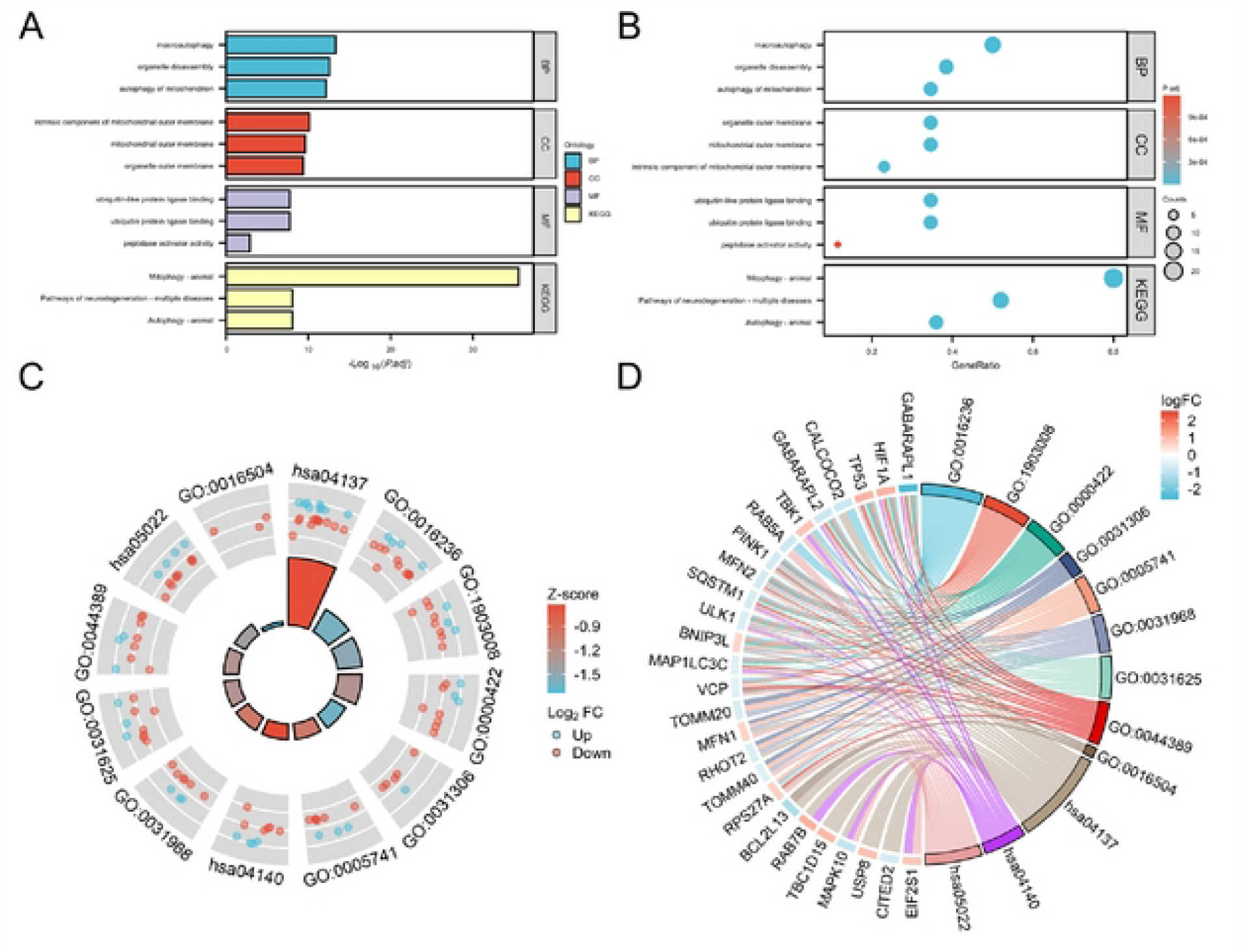
GO enrichment analysis of 27 differentially expressed mitophagy-related genes, including BPs, CCs and MFs. (A). Bar plot of enriched GO terms. (B).Bubble plot of enriched GO terms.(C). Eight Diagrams of enriched GO terms. GO. Gene Ontology; BPs. biological processes; CCs. cellular components; MFs, molecular functions; DEGs, differentially expressed genes. (D). Chordal graph of enriched GO terms It shows the relationship between DEGs and the first 9 enriched GO pathways.

**FIGURE 5.**
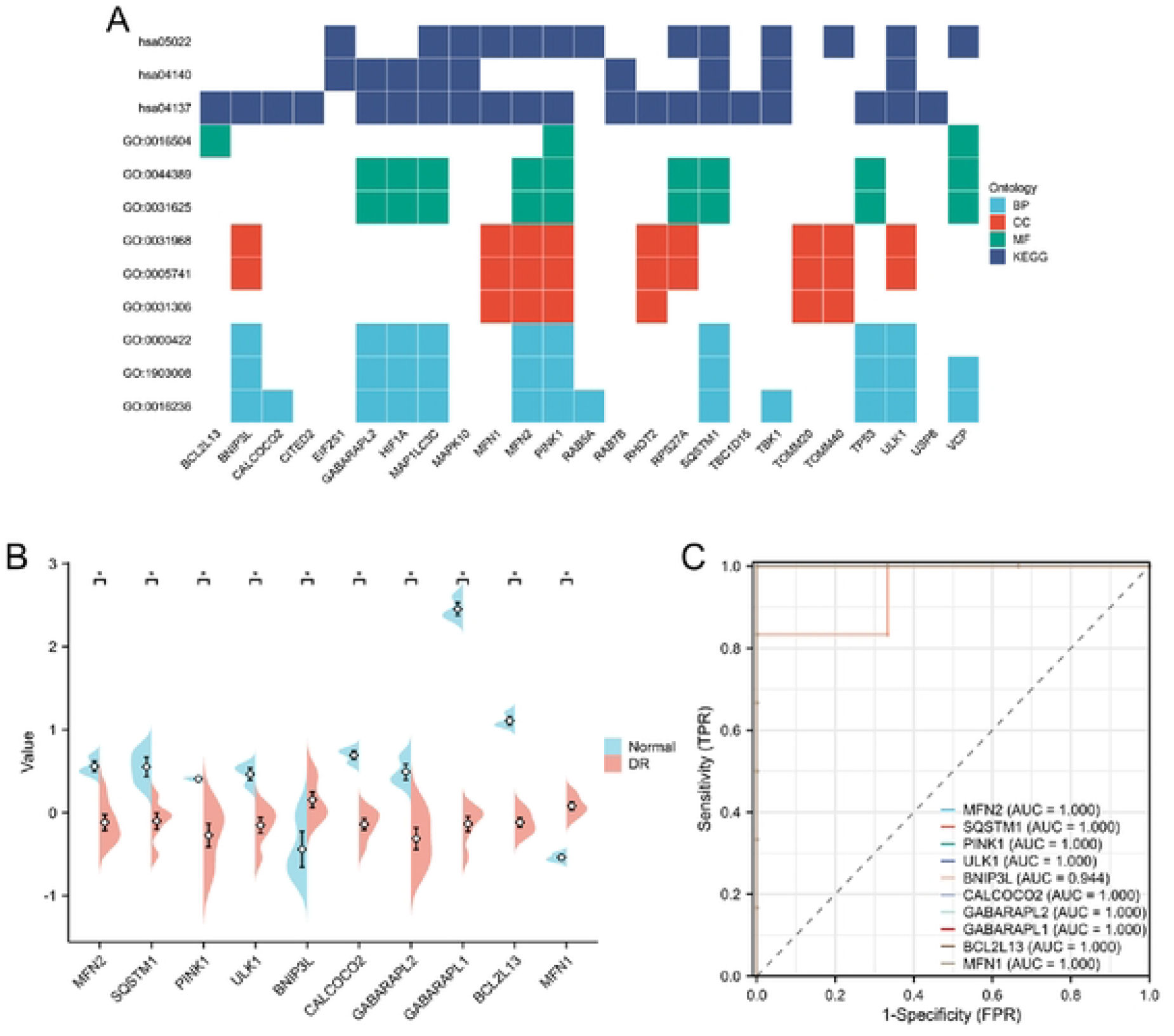
(A). Heatmap-like functional classification. (B). The bean plot shows expresion differences of 10 hub genes in dr and control groups. (C),ROC curve of 10 hub genes

### 3.4. PPI Network Analysis and Hub Gene Identification

We imported 27 differentially expressed mitophagy genes into STRING to construct a protein-protein interaction network.Selected 10 hub genes with the highest value using Cytoscape software (v3.9.0). BNIP3L,MFN1 genes were up-regulated, and GABARAPL1,BCL2L13,CALCOCO2,GABARAPL2,PINK1,MFN2,SQSTM1,ULK 1 were down-regulated.Impaired mitophagy function may be closely related to the occurrence and development of DR. (Figures 6A, B,Table 1)

**Table 1.**
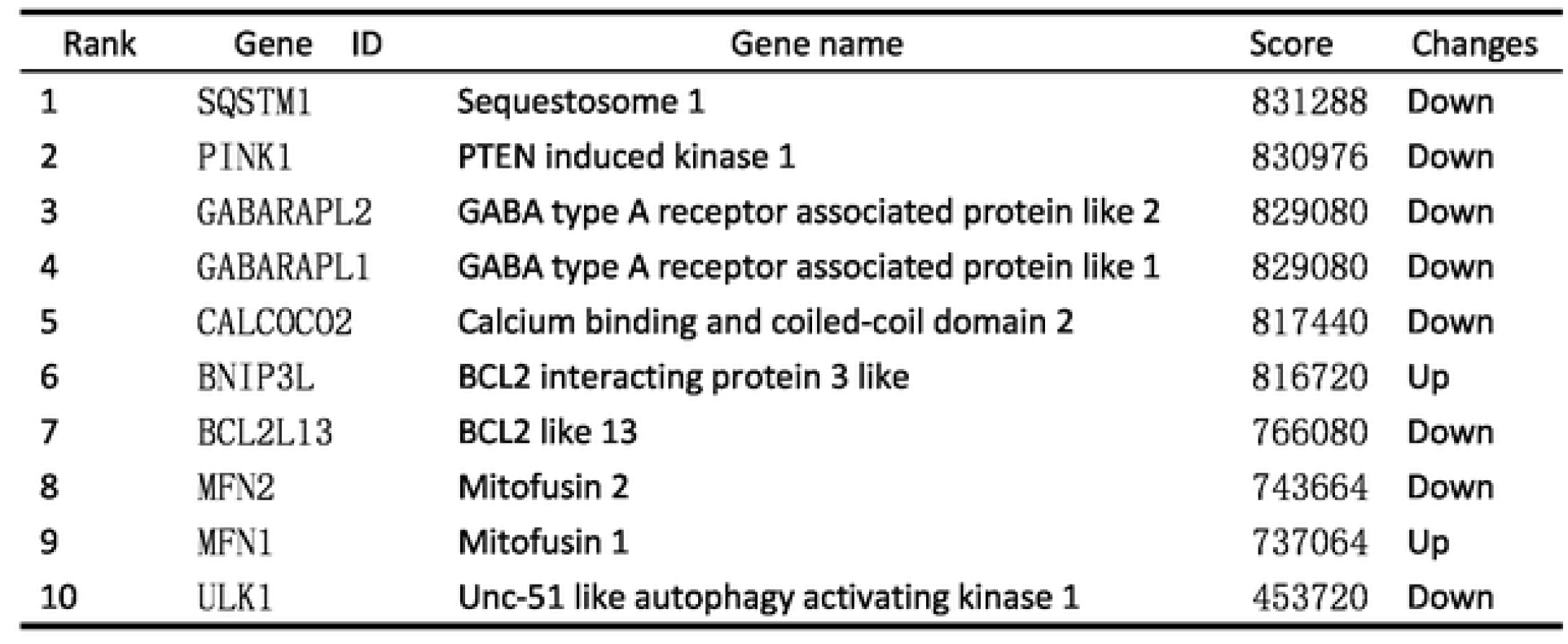
Top 10 in network ranked by Mcc method.

**FIGURE 6.**
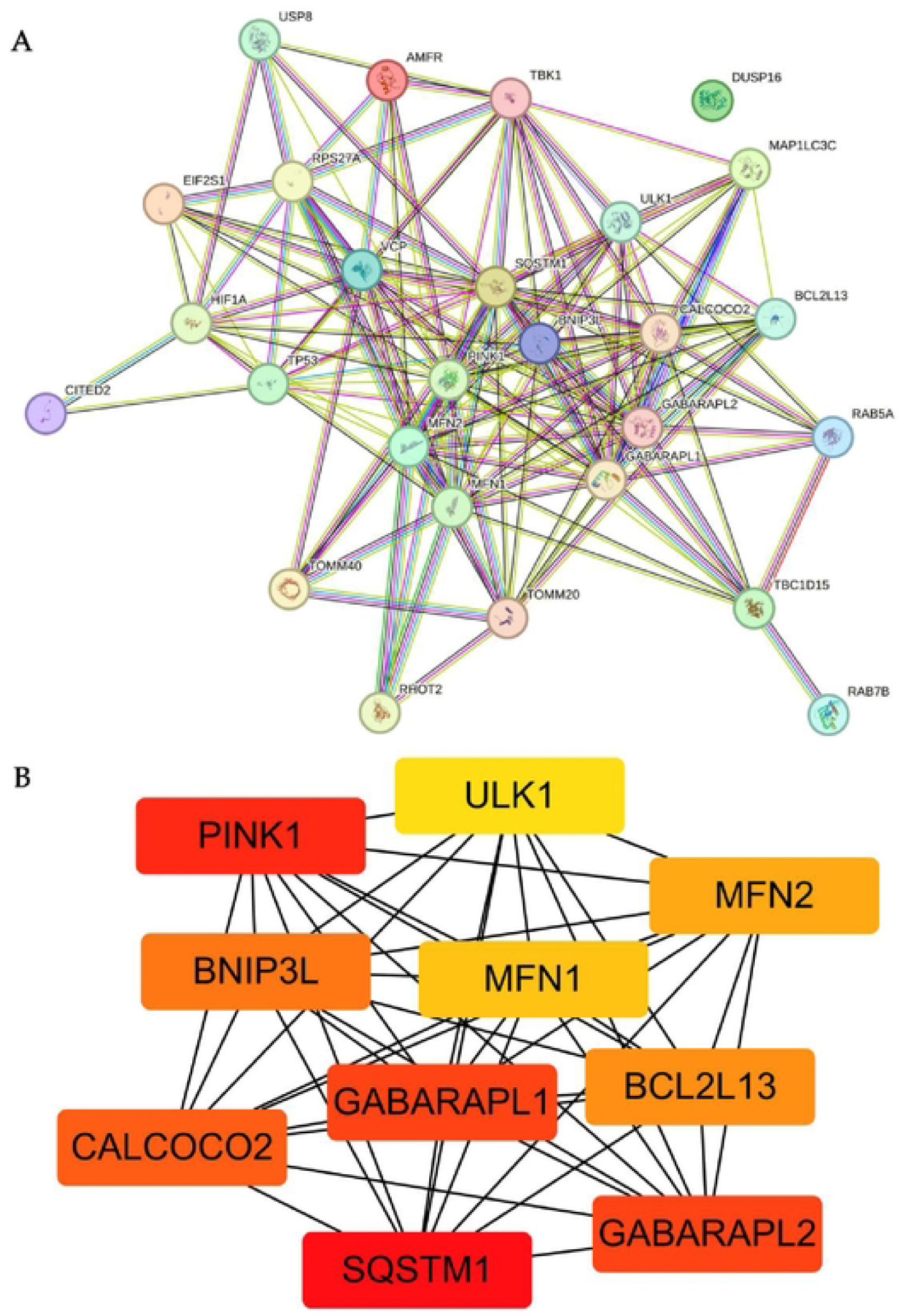
Construction of PPI network and identification of hub genes (A), The PPI between 27 differentially expressed mitophagy-related genes was constructed byusing the STRING database. The node represents the gene, and the edge represents the relationship between the genes. (B). The top 10 key genes were screened through the PPI network map. Different colors in the image only represent different genes and have no other substantive meaning PPI. protein-protein interaction.

### 3.5. Validation of differentially expressed mitophagy-related genes in a diabetes model

In diabetic retinopathy (DR), there is impairment in the mitophagy of retinal microvascular endothelial cells, leading to impaired mitochondrial function and the occurrence of adverse physiological reactions such as oxidative stress^[9]^.Human retinal microvascular endothelial cells were cultured in high glucose (30mM) to simulate a diabetic retinopathy model in vitro.In the qRT-PCR validation results, the gene expression of PINK1 and GABARAPL1 is downregulated, while the gene expression of BNIP3L and MFN1 is upregulated in HG treated HRMEC cells compared with the normal control cells,the expression results of these genes are consistent with our bioinformatics analysis.In the qRT-PCR validation results, there were no significant differences in the expression of SQSTM1,ULK1,BCL2L13,CALCOCO2,GABARAPL2,MFN2 between the HG and control groups(Figures 7).we studied the differential expression of PINK1,BNIP3L and MFN1, the markers of mitophagy, among NG group and HG group by western blot.The results revealed that the protein expression levels of PINK1 and MFN1 were downregulated in HRCECs under the high-glucose 48h enviroment and the protein expression level of BNIP3L was upregulated in HRCECs under the 48h high-glucose enviroment.

**FIGURE 7.**
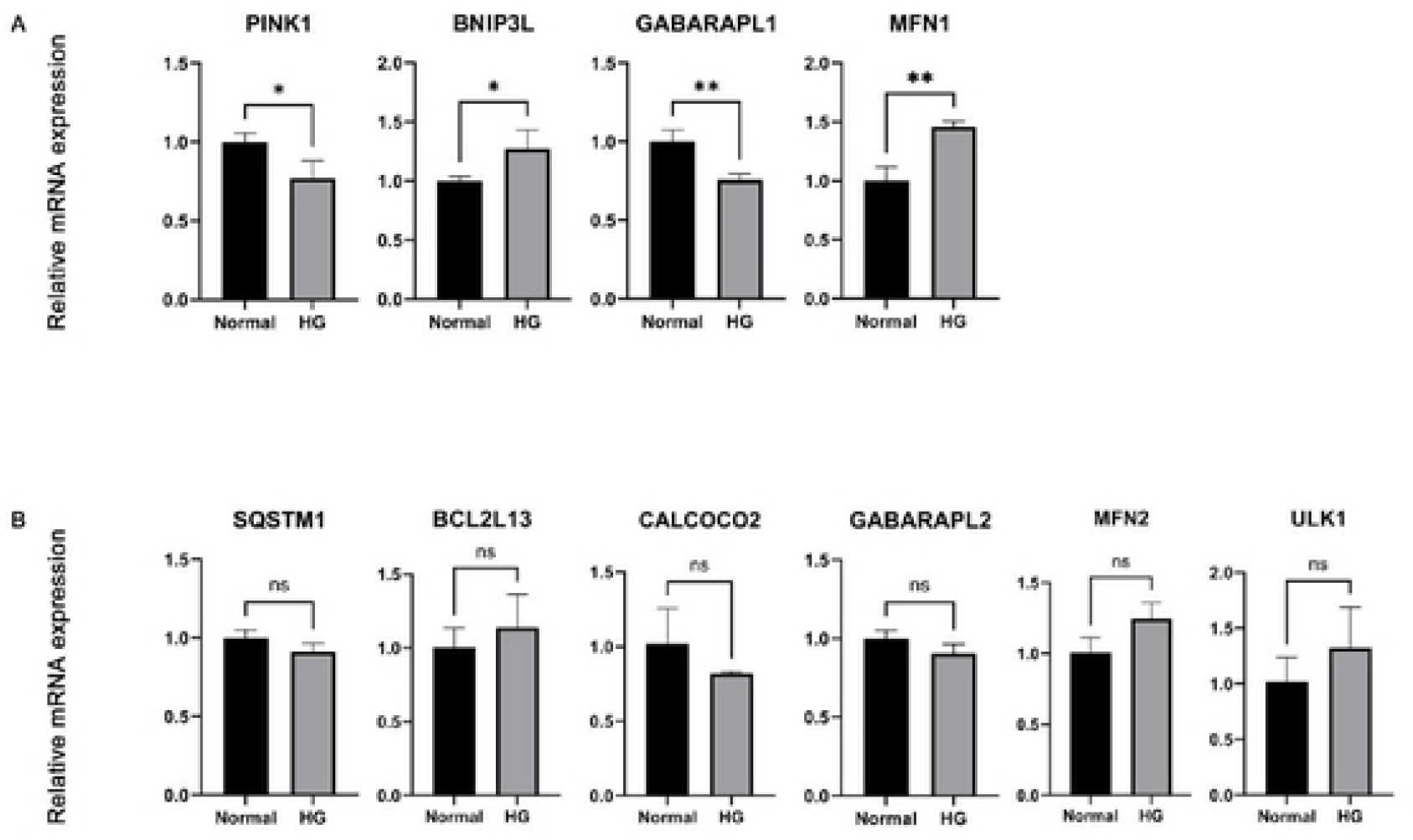
The mRNA level of 10 hub genes were measured in HRCECs cels. (A). The mRNA level of PINK1. BNIP3L, GABARAPL1 and MFN1 were evaluated in cells amples by qRT-PCR (B). The mRNA level of SQSTM1, BCL2L13, CALCOCO2.GABARAPL2. MFN2 and ULK1 were measured In cell samples by qRT-PCR. P-values were calculated using a two-sided unpaired Student s l-test. *P < 0.05; **P < 0.01; ns, non-significant. HRCECs, human retinal microvascular endothelial cells; qRT-PCR. quantitative real-time polymerase chain reaction.

**FIGURE 8.**
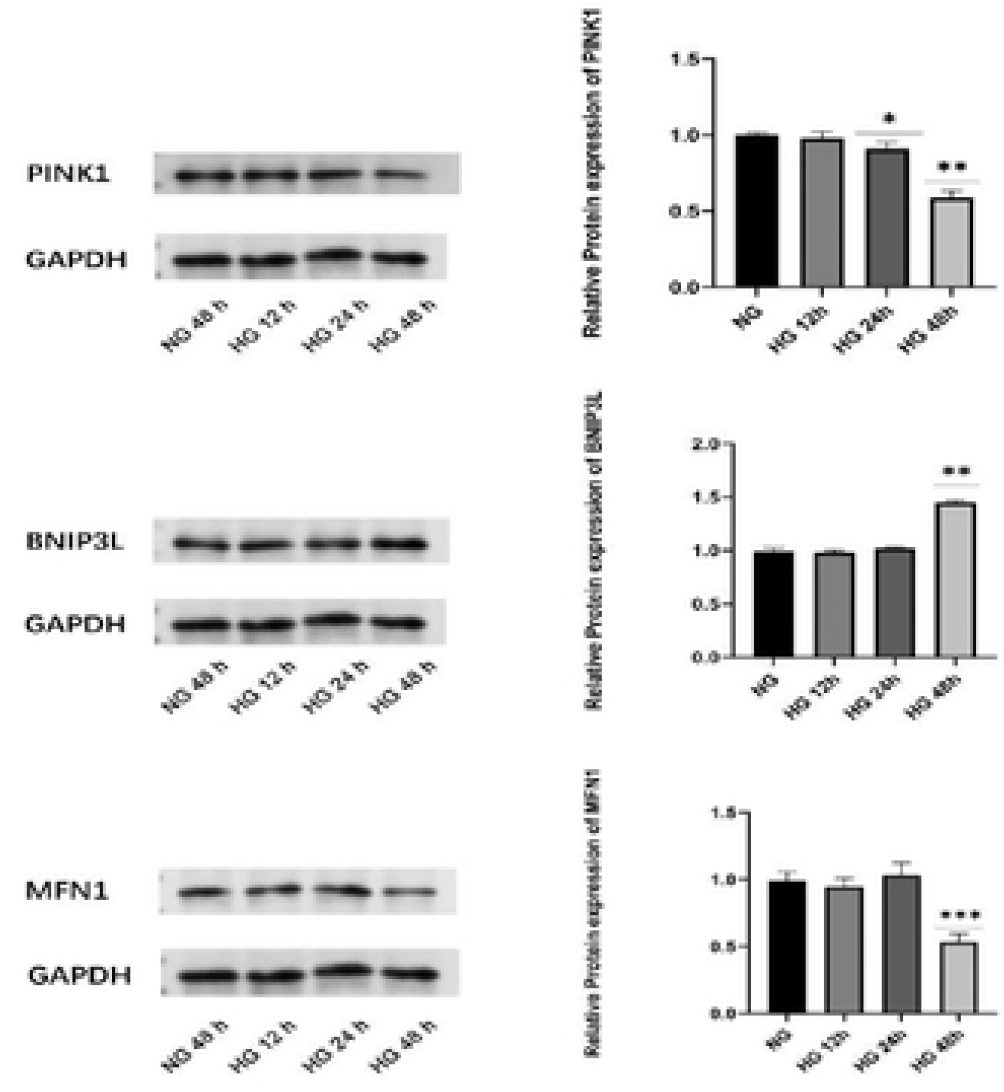
The protein levels of PINK1, BNIP3L and MFN1 were evaluated in cell samples by western blot.

## 4. Discussion

Diabetic retinopathy is a common microvascular complication of diabetes and a leading cause of blindness worldwide^[12]^. Despite the three-tiered treatments for DR (anti-VEGF therapy, retinal laser photocoagulation, and vitrectomy) being able to delay disease progression, DR remains incurable at present^[13]^.Therefore, uncovering the pathogenesis of DR is crucial for its treatment and prognosis.High glucose environment can trigger a series of molecular, biochemical, and functional abnormalities, affecting many metabolic pathways^[14]^, including the activation of protein kinase C, increased advanced glycation end products (AGEs)^[15]^, and the polyol and hexosamine pathways. It has been reported that high glucose significantly inhibits the proliferation and migration of human retinal microvascular endothelial cells(HRCECs)^[16]^ and induces abnormal synthesis of extracellular matrix, leading to basal membrane thickening^[17]^, which is a typical early characteristic of DR.

Mitophagy, first proposed by Lemasters in 2005, is a selective process in which cells remove damaged or dysfunctional mitochondria in response to stimuli, thereby preserving functional mitochondrial populations. It is considered a major mechanism for controlling the number and quality of mitochondria^[18]^. Mitophagy dysfunction can lead to mitochondrial accumulation, changes in mitochondrial cristae structure, bio energetic dysfunction, cell death, and ultimately, potentially fatal organ and tissue damage^[19]^.The functionality of mitochondria was compromised after 1 day of exposure to high glucose, but improved after 10 days of exposure to high glucose, and this change required an increase in the clearance of damaged mitochondria(mitophagy).The results indicate that high glucose-induced mitochondrial adaptation may be a contributing factor to the delayed onset of DR, while the loss of mitochondrial adaptation including mitophagy capability may set the stage for the development of DR^[20]^.During DR, the level of mitophagy upregulation or inhibition may depend on the concentration of high glucose.Zhang^[21]^ observed a slight increase in glucose concentration (15 mmol/L) inducing upregulation of mitophagy in cultured retinal pigment epithelial cells, while a substantial increase in glucose concentration (50 mmol/L) inhibited mitophagy and led to apoptotic cell death.The mechanism may involve mild high glucose-induced stress response, prompting cells to engage in mitophagy for “self-protection,” while severe or sustained high glucose levels can cause cellular damage, leading to impairment of mitophagy. Therefore, understanding the molecular mechanisms involved in DR is of significant clinical importance for effective intervention in the disease.However, research on mitophagy in the field of retinal diseases is still limited, and the underlying mechanisms remain unclear, with their exact roles not well defined. Therefore, based on bioinformatics analysis, we explore the potential pathogenic mechanisms of mitophagy in the occurrence and development of diabetic retinopathy (DR). Elucidating the potential relationship between mitophagy and DR may identify new diagnostic biomarkers and provide pharmacological support for targeted therapies, offering new avenues for treatment strategies targeting mitophagy In this study,we initially used bioinformatics analysis to identifiy 27 potential mitophagy genes related with diabetic retinopathy.Next, we further identified 10 hub genes related to DR,including BNIP3L,MFN1,GABARAPL1,BCL2L13,CALCOCO2,ULK1 PINK1,MFN2,SQSTM1 by using PPI network. The function of these genes in the occurrence of DR has been extensively studied. For example,PTEN-induced putative kinase 1 (PINK1)/Parkin mitophagy pathway is one of the most characteristic ubiquitin-dependent pathways, mediated by PINK1 and Parkin proteins^[19]^.However, when mitochondrial function is impaired, PINK1 degradation is inhibited and it accumulates, leading to phosphorylation and recruitment to activate Parkin protein in the cytoplasm^[22-23]^.Activated Parkin protein binds to damaged mitochondria and ubiquitinates mitochondrial outer membrane proteins, thereby recruiting the autophagy receptor p62/SQSTM1 (sequestosome 1, referred to as p62), which can bind to the microtubule-associated protein 1 light chain 3 (microtubule-associated protein 1A/1B-light chain 3, LC3) on the autophagosome membrane, promoting the engulfment of mitochondria for autophagic degradation, completing mitophagy^[24]^.BNIP3 and Nix/BNIP3L share 50% homology and are associated with autophagy, particularly in relation to mitophagy induced by hypoxia ^[25]^. Under hypoxic conditions, Nix and BNIP3 are regulated to upregulate their expression through hypoxia-inducible factor 1α (HIF-1α) ^[26]^. Studies have identified that the structure of Nix/BNIP3L includes a single-pass carboxy-terminal transmembrane domain that targets the outer mitochondrial membrane (OMM), and it contains an LC3-interacting region (LIR). Thus, Nix/BNIP3L can directly interact with LC3/GABARAP-related proteins to induce specific mitophagy^[27]^.Studies have shown that GABARAPL1 can bind to mitophagosomes and participate in the process of mitophagy. Additionally, it is involved in the glycolytic process, can relocate to damaged mitochondria, and interacts with BNIP3L. However, the specific mechanisms by which it participates in the mitophagy process are not yet fully understood^[28]^.MFN1 is involved in ensuring the fusion function of the mitochondrial outer membrane^[29]^. Mitochondrial fusion has a protective role under physiological conditions, but its role in ischemia-reperfusion injury remains controversial^[30]^. Research has found that knocking out the MFN1 gene provides protection against myocardial infarction, with mechanisms related to alleviating oxidative stress^[31]^.

## 5. Conculsions

Among 10 predicted DR related hub genes, we showed that the expression of MFN1,BNIP3L,GABARAPL1 and PINK1 were consistent with that of bioinformatics analysis of mRNA chip.These genes are potential novel biomarkers for the diagnosis of DR and its targeted therapy. To sum up, our study contributes new insights into the pathogenesis associated with mitophagy, as well a theoretical basis for discovering new diagnostic markers and therapeutic methods in DR.

## Notes

### Competing Interest Statement

The authors have declared no competing interest.

